# Population-scale dietary signatures of the oral microbiome

**DOI:** 10.64898/2026.10.28.762516

**Authors:** Kaiyu Liang, Haochen Xue, Anastasia Godneva, Jionglong Su, Xiaojun Zhu, Huifa Li, Yulong Li, Sibei Chen, Yichen Li, Eran Segal, Imran Razzak

## Abstract

Diet is an environmental factor that influences oral microbial communities. How specific foods, nutrients and dietary patterns relate to microbial diversity, species composition and functional potential remains incompletely characterised. Here, we combined detailed dietary records with buccal metagenomic profiles from 5,824 Human Phenotype Project participants. Using these data, we examined diet–oral microbiome associations with prediction models and covariate-adjusted analyses. Plant-based and guideline-based dietary patterns showed similar associations across oral microbial species. However, species within the same genus could show opposite associations with a dietary pattern. For example, a whole-food plant-based diet score was positively associated with *Streptococcus sanguinis* but negatively with *S. gordonii* and *S. oralis*. Some diet–oral microbiome associations were replicated at the genus level in an independent US National Health and Nutrition Examination Survey cohort of 4,667 participants. We map dietary associations with oral microbial diversity, species composition and functional potential, revealing shared microbial signatures across dietary patterns.

## 1 Introduction

The oral microbiome is repeatedly exposed to dietary components as food and drink enter the mouth [1]. Compared with the gut microbiome, the oral microbiome is more directly and repeatedly exposed to food residues, fermentable carbohydrates, acidic beverages, dietary nitrate, polyphenols, dairy products and fermented foods [2, 3]. This direct exposure makes the oral cavity a unique ecological interface for studying how habitual diet relates to the human microbiome.

Diet may shape the oral microbiome through multiple ecological and metabolic mechanisms. Frequent intake of fermentable sugars and refined carbohydrates can enrich acidogenic and aciduric taxa, lower dental plaque pH and promote caries-associated dysbiosis [4–6]. In contrast, diets rich in fruits, vegetables and plant-derived bioactive compounds may help maintain a more stable oral microbial ecosystem and influence functional microbial groups, including taxa involved in nitrate metabolism [7–9]. Dairy products and probiotic-containing foods may also influence the oral ecosystem by altering pH, calcium–phosphate balance, microbial colonization or the transfer of food-associated microbes [10–12].

Previous population-based studies have linked specific dietary components and overall diet quality to the oral microbiome [3, 13–15]. However, such dietary assessment often relies on food-frequency questionnaires, which estimate usual intake [3, 13, 14]. These provide limited information on foods consumed at individual eating occasions, intake timing and variation across days. Oral microbiome profiling often uses 16S rRNA gene sequencing, which provides limited species-level resolution. This approach also does not directly measure microbial functional gene content [16]. Population-scale studies combining detailed, multi-day dietary records with species-level and functional oral metagenomic profiles remain limited. Consequently, how individual foods, nutrients and dietary patterns relate to oral microbial species and functional potential remains incompletely characterised.

To address these gaps, we leveraged the Human Phenotype Project (HPP). From the broader HPP cohort, we analysed 5,824 participants who had matched app-based dietary logs, oral shotgun metagenomic profiles, and covariate data associated with the same baseline visit, including age, sex, body mass index (BMI), smoking status, and oral-health variables. The app-based dietary logs covered multiple days around the baseline visit, capturing individual foods, portion sizes, intake times and corresponding nutritional information. These records allowed us to quantify diet at several levels, including individual foods, nutrients, food categories and dietary patterns, while also describing eating behaviours and day-to-day variation in intake. In addition, the dataset included paired shotgun metagenomic profiles of the oral microbiome, enabling both species-level and functional pathway characterization [16].

Based on this paired dietary and metagenomic dataset, we systematically examined how diet relates to the oral microbiome across multiple biological levels. We trained multivariable machine-learning models to predict oral microbial alpha diversity, species-level relative abundances and functional pathways from dietary features. We then used feature-attribution analyses to identify the individual foods, nutrients and dietary patterns contributing to these predictions. Together, this study provides a population-scale, high-resolution map of diet–oral microbiome associations, linking detailed dietary intake to oral microbial composition and functional potential.

## 2 Results

### 2.1 Study cohort and research design

We assembled a matched diet–oral microbiome dataset from the Human Phenotype Project (HPP), a deeply phenotyped population cohort with app-based dietary records and shotgun metagenomic profiling of oral samples (Fig. 1a). Detailed sample collection and metagenomic processing procedures are described in Methods 4.3.

**Fig. 1.**
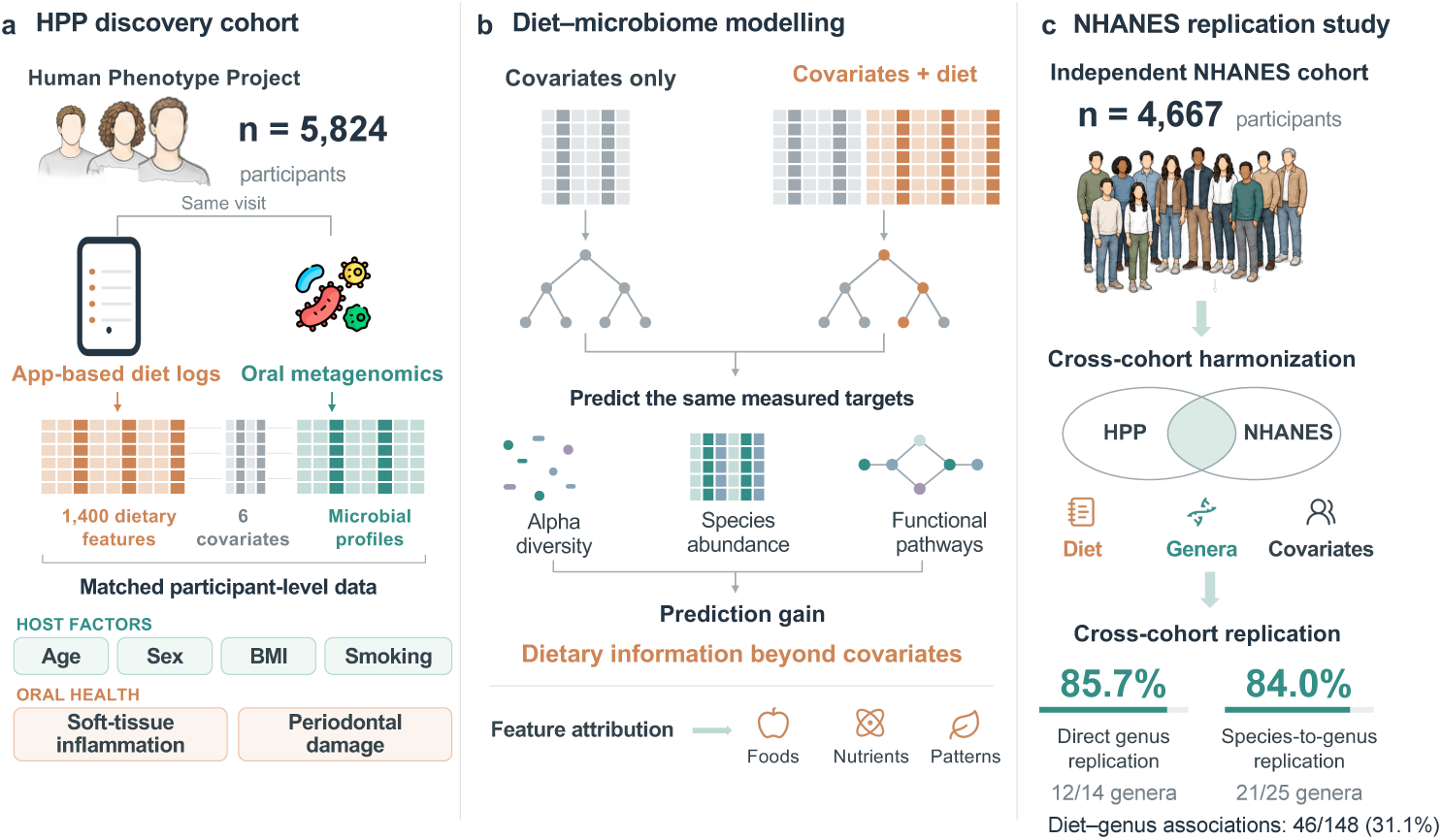
Study design for diet–oral microbiome discovery and cross-cohort replication. **a**, App-based dietary records yielded 1,400 features and were paired with oral metagenomic profiles from 5,824 Human Phenotype Project (HPP) participants at the same baseline visit. **b**, Covariates-only and covariates-plus-diet models were compared for predicting alpha diversity, species abundance and functional pathways. Feature attribution identified contributing foods, nutrients and dietary patterns. **c**, Genus-level replication in 4,667 National Health and Nutrition Examination Survey (NHANES) participants, with 12 of 14 genera replicated in the direct genus-level analysis and 21 of 25 in the species-to-genus analysis. Individual diet–genus associations replicated for 46 of 148 HPP candidate pairs. Replication criteria are described in Methods 4.7.

Dietary intake was recorded using the HPP diet-logging smartphone app. Participants were asked to begin logging one day before their baseline HPP visit and to record everything they ate and drank for at least 14 days afterwards. The app remained available throughout the study. Each logged entry included consumption time, food identity, serving amount, food category and nutritional information, including calories, macronutrients, sodium, alcohol and dietary fibre. Meals were defined as clock hours containing at least one logged entry.

From these logs, we generated dietary features spanning multiple levels of intake: individual foods, food categories, nutrients, macronutrient composition, eating timing, logging behaviour and diet-quality or dietary-pattern scores (Fig. 1a; Methods 4.2). These included measures such as total energy intake, food diversity, meals per day, foods per day, fasting window, eating-window duration, fibre intake, macronutrient calorie percentages and diet-pattern scores including whole-food plant-based, EAT– Lancet, Mediterranean-like and other dietary adherence scores.

Matching dietary records, oral metagenomic profiles and host covariates from the same baseline HPP visit yielded a dataset of 5,824 participants. Age and summary dietary-recording and intake characteristics stratified by sex appear in Table 1. Using this matched visit-level dataset, we tested whether dietary profiles were associated with oral microbiome structure and function beyond baseline covariates (Fig. 1b). We evaluated three microbiome target layers: oral alpha diversity, species-level relative abundance and functional pathway abundance. For each target, we compared covariates-only models with covariates-plus-diet models. Both model types included covariates associated with the same baseline visit: age, sex, body mass index, smoking status, soft-tissue inflammation level and periodontal damage level (Methods 4.1). We assessed prediction using out-of-fold Pearson correlation, evaluated significance with permutation testing and false discovery rate (FDR) correction, and used SHapley Additive exPlanations (SHAP) to identify dietary and host features contributing to model predictions.

**Table 1.**
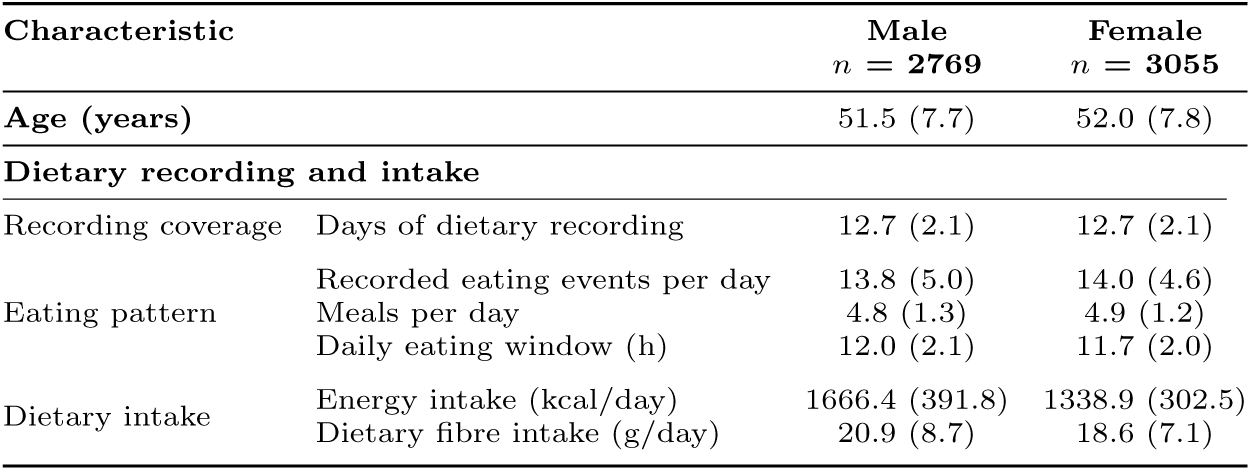
Baseline characteristics of the matched diet–oral microbiome cohort stratified by sex. Values are mean (s.d.).

### 2.2 Diet and oral alpha diversity: associations and prediction

We tested whether baseline dietary features improved out-of-fold (OOF) prediction of oral alpha diversity beyond age, sex, BMI, smoking and oral-health covariates. We evaluated species richness and Shannon diversity using covariates-only and covariates-plus-diet models.

Adding dietary features increased OOF Pearson’s *r* from 0.078 to 0.145 for species richness (Δ*r* = 0.067) and from 0.066 to 0.121 for Shannon diversity (Δ*r* = 0.055; Fig. 2a). Corresponding *R*^2^ values increased from 0.005 to 0.020 and from 0.003 to 0.014, respectively, indicating improved prediction after adding dietary features. Covariates-plus-diet models significantly predicted species richness and Shannon diversity after permutation testing and FDR correction (permutation *P* = 0.004; FDR *q* = 0.004 for each outcome).

**Fig. 2.**
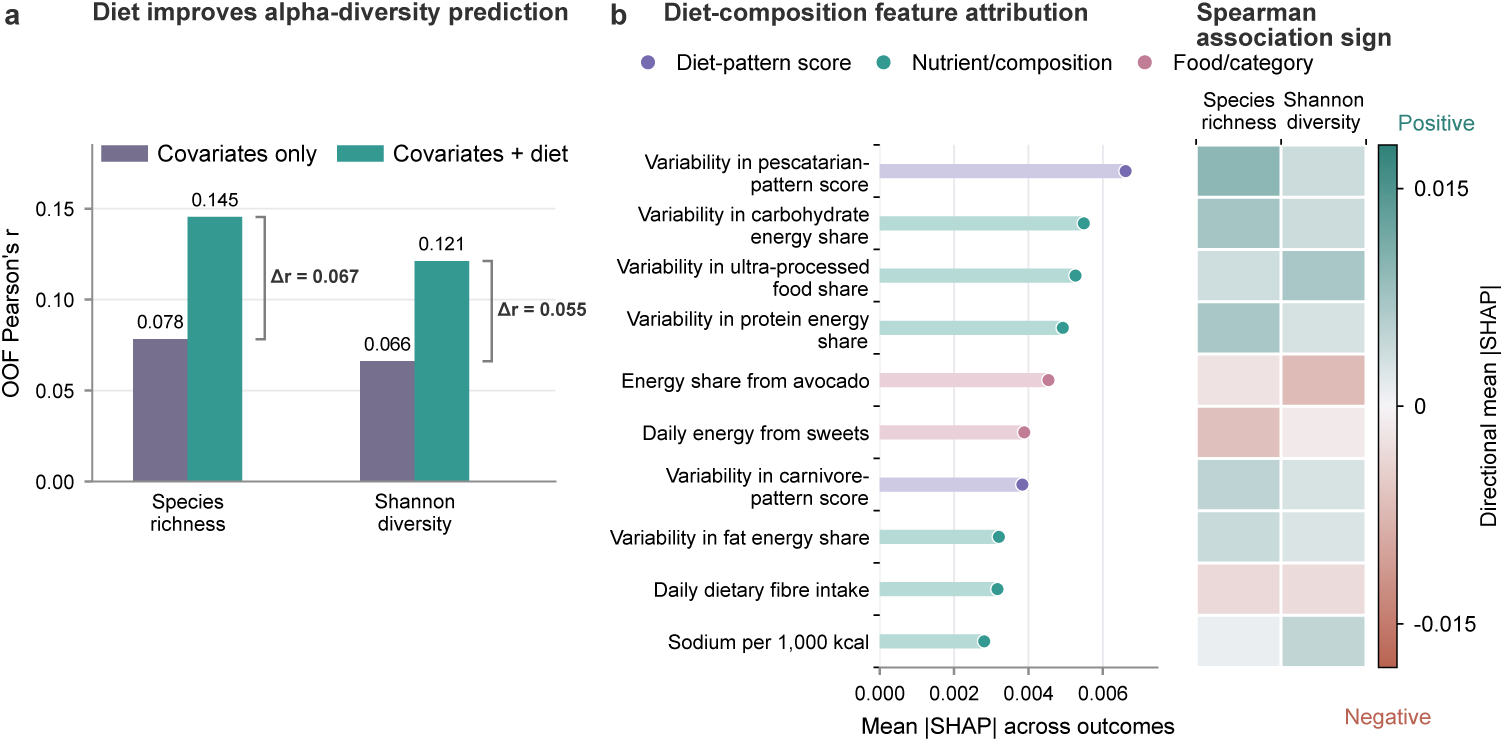
Dietary features improve prediction of oral alpha diversity. **a**, Out-of-fold (OOF) Pearson’s *r* for covariates-only and covariates-plus-diet models predicting species richness and Shannon diversity. Brackets show the increase in *r* after adding dietary features (Δ*r*). **b**, Bars show mean absolute SHAP values for ten selected dietary-composition features, averaged across the two outcomes. Heatmap colours indicate mean absolute SHAP values for each outcome, with signs determined by feature–outcome Spearman correlations.

We then used SHAP-based feature attribution to identify features contributing to alpha-diversity prediction. Age had the highest mean absolute SHAP value averaged across the two outcomes. The daily numbers of recorded eating events, food categories, meals and food items also ranked among the leading predictors. Fig. 2b shows ten selected dietary-composition features. Among these, day-to-day variability in pescatarian-pattern score had the highest mean absolute SHAP value across the two outcomes. This was followed by variability in carbohydrate energy share, ultra-processed food share and protein energy share. Food-level contributors included the energy share from avocado and daily energy from sweets.

Both diversity outcomes shared several directional profiles but differed in magnitude (Fig. 2b). Day-to-day variability in pescatarian- and carnivore-pattern scores was positively associated with both outcomes. This direction was observed for variability in macronutrient energy shares and ultra-processed food share. By contrast, energy share from avocado, daily energy from sweets and daily dietary fibre intake had negative directions for both outcomes. The negative association between sweets and richness is consistent with lower oral richness reported with high sucrose intake and high-sugar beverage consumption [6, 17, 18]. Sodium intake per 1,000 kcal had greater attribution magnitude for Shannon diversity than for species richness.

Covariate-adjusted analyses of mean dietary-pattern scores showed that vegan, vegetarian and pescatarian scores were negatively associated with both species richness and Shannon diversity. AHEI, DASH and EAT–Lancet scores showed the same negative associations. The low-carbohydrate score was positively associated with both outcomes. Whole-food plant-based and carnivore scores were negatively and positively associated with species richness, respectively. These associations were significant in both the primary model and the model additionally adjusted for oral-health variables (FDR *q <* 0.05).

### 2.3 Prediction of oral species-level abundance from diet

We extended the prediction analysis from community diversity to the abundances of individual oral species. After prevalence filtering, 136 oral species were retained for modelling (Methods 4.3). For each species, we compared covariates-only and covariates-plus-diet models (Methods 4.5; Fig. 3).

**Fig. 3.**
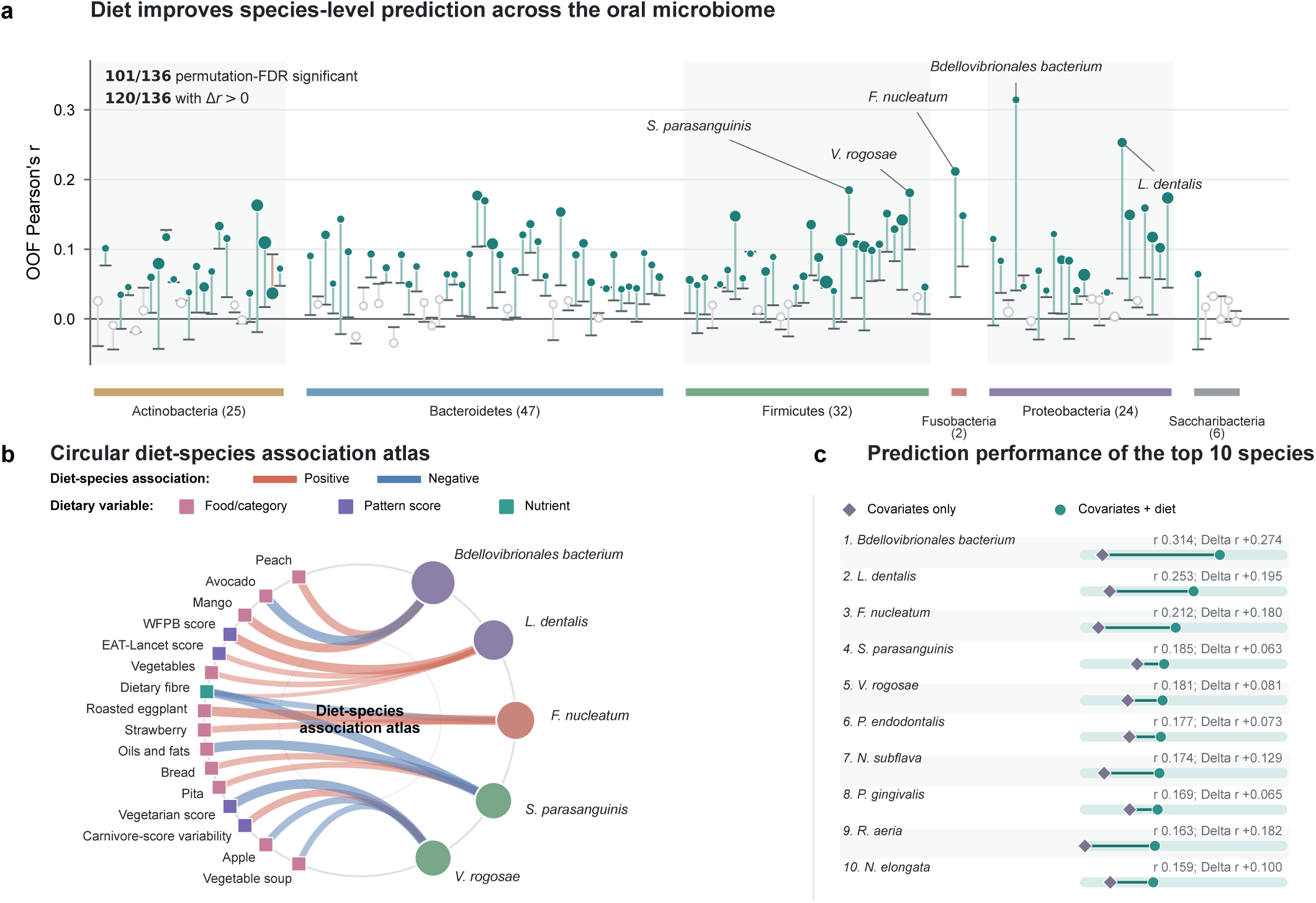
Dietary features predict oral species-level abundance. **a**, Out-of-fold (OOF) prediction performance across 136 retained species grouped by phylum. Horizontal dashes and circles show Pearson’s *r* for covariates-only and covariates-plus-diet models, respectively. Filled circles indicate permutation-FDR-significant species; circle area represents species prevalence. **b**, Diet–species attribution atlas for the five species with the highest covariates-plus-diet OOF Pearson’s *r*. Link colour indicates the direction of the feature–species Spearman correlation. Link width represents mean absolute SHAP normalized to the largest displayed value within each species. Species-node colour denotes phylum, and node area represents covariates-plus-diet OOF Pearson’s *r*. WFPB, whole-food plant-based. **c**, Prediction performance of the ten highest-ranked species, ordered by covariates-plus-diet OOF Pearson’s *r*. Abbreviated species names include *S. parasanguinis*, *Streptococcus parasanguinis*; *V. rogosae*, *Veillonella rogosae*; and *P. endodontalis*, *Porphyromonas endodontalis*. *N. subflava*, *P. gingivalis*, *R. aeria* and *N. elongata* denote *Neisseria subflava*, *Porphyromonas gingivalis*, *Rothia aeria* and *Neisseria elongata*, respectively.

Covariates-plus-diet models significantly predicted 101 of 136 oral species (74.3%) after permutation-based FDR correction (Fig. 3a; Supplementary Fig. 1). Prediction correlations varied across species, with a median Pearson’s *r* of 0.063 (Fig. 3a). The strongest examples included Bdellovibrionales bacterium (*r* = 0.314), *Lautropia dentalis* (*r* = 0.253), *Fusobacterium nucleatum* (*r* = 0.212) and *Streptococcus parasanguinis* (*r* = 0.185) (Fig. 3c). The median improvement was Δ*r* = 0.043 (interquartile range, 0.014–0.073), with 120 of 136 species showing a positive Δ*r* relative to the covariates-only baseline (Fig. 3a). Selected species showed higher OOF prediction correlations than either alpha-diversity outcome.

Across significant species, age and BMI were among the strongest background predictors. Population studies have also linked oral microbial composition to age and adiposity [19, 20]. Eating-pattern and dietary-recording features also contributed broadly, including the average number of recorded eating events, fasting window length, daily meal frequency, eating window duration and daily food-item diversity. Diet-related predictors included dietary fibre, fruit intake, vegetable intake, coffee, avocado, watermelon and diet-quality scores, suggesting that species-level oral microbiome profiles captured both specific food exposures and broader dietary structure.

Dietary attribution profiles differed among the five best-predicted species (Fig. 3b). Bdellovibrionales bacterium showed prominent fruit-related signals involving avocado, mango and peach. *Lautropia dentalis* showed a more plant-forward profile, with contributions from whole-food plant-based score, EAT–Lancet score, vegetable intake and dietary fibre. In patients undergoing endoscopy, greater fruit and vegetable intake was associated with higher salivary *Lautropia* sp. abundance [21]. *Fusobacterium nucleatum* showed contributions from roasted eggplant, strawberry and dietary fibre. For *Streptococcus parasanguinis*, leading dietary attributions involved oils and fats, dietary fibre, bread and pita, whereas those for *Veillonella rogosae* involved vegetarian-pattern score, day-to-day variability in carnivore-pattern score, apple and vegetable soup. After adjustment for oral sampling season, dietary prediction gains (Δ*r*) remained positive for all five species.

### 2.4 Dietary patterns show shared and species-specific oral microbial association signatures

To compare microbial association profiles across dietary patterns, we tested 1,768 associations between 13 dietary pattern scores and 136 retained species. The primary covariate-adjusted model identified 686 FDR-significant associations. After additional oral-health adjustment, 653 associations were significant in both models, all with concordant directions.

The number and direction of associated species differed substantially across dietary patterns. Plant-forward and guideline-based scores were dominated by negative associations. The EAT–Lancet score was associated with 75 species, including 61 negative associations; the Alternative Healthy Eating Index (AHEI) score with 69 species, including 56 negative associations; and the whole-food plant-based score with 66 species, including 50 negative associations. In contrast, carnivore and low-carbohydrate scores were dominated by positive associations. Of the associations identified for the carnivore, low-carbohydrate and extremely low-carbohydrate scores, 33 of 39, 38 of 48 and 24 of 30 were positive, respectively.

Across the 13 dietary patterns, 16 of 26 associations with species richness or Shannon diversity were FDR significant in both adjusted models. The inverse association between AHEI and richness was consistent with lower observed subgingival richness at higher AHEI scores in ORIGINS [22].

Across the 136 retained species, the dietary patterns showed a clear structure of shared and opposing association profiles (Fig. 4a,b). All 36 pairwise correlations among the nine plant-forward or guideline-based scores were positive, with a median Spearman *ρ* of 0.803 (interquartile range, 0.678–0.929). The carnivore, low-carbohydrate and extremely low-carbohydrate scores also showed similar profiles, with a median pairwise *ρ* of 0.712. Ultra-processed food share had equal numbers of positive and negative species associations (14 each). Across all 136 retained species, however, its association profile was negatively correlated with those of 11 of the 12 other patterns (median Spearman *ρ* = *−*0.541; interquartile range, *−*0.600 to *−*0.443).

**Fig. 4.**
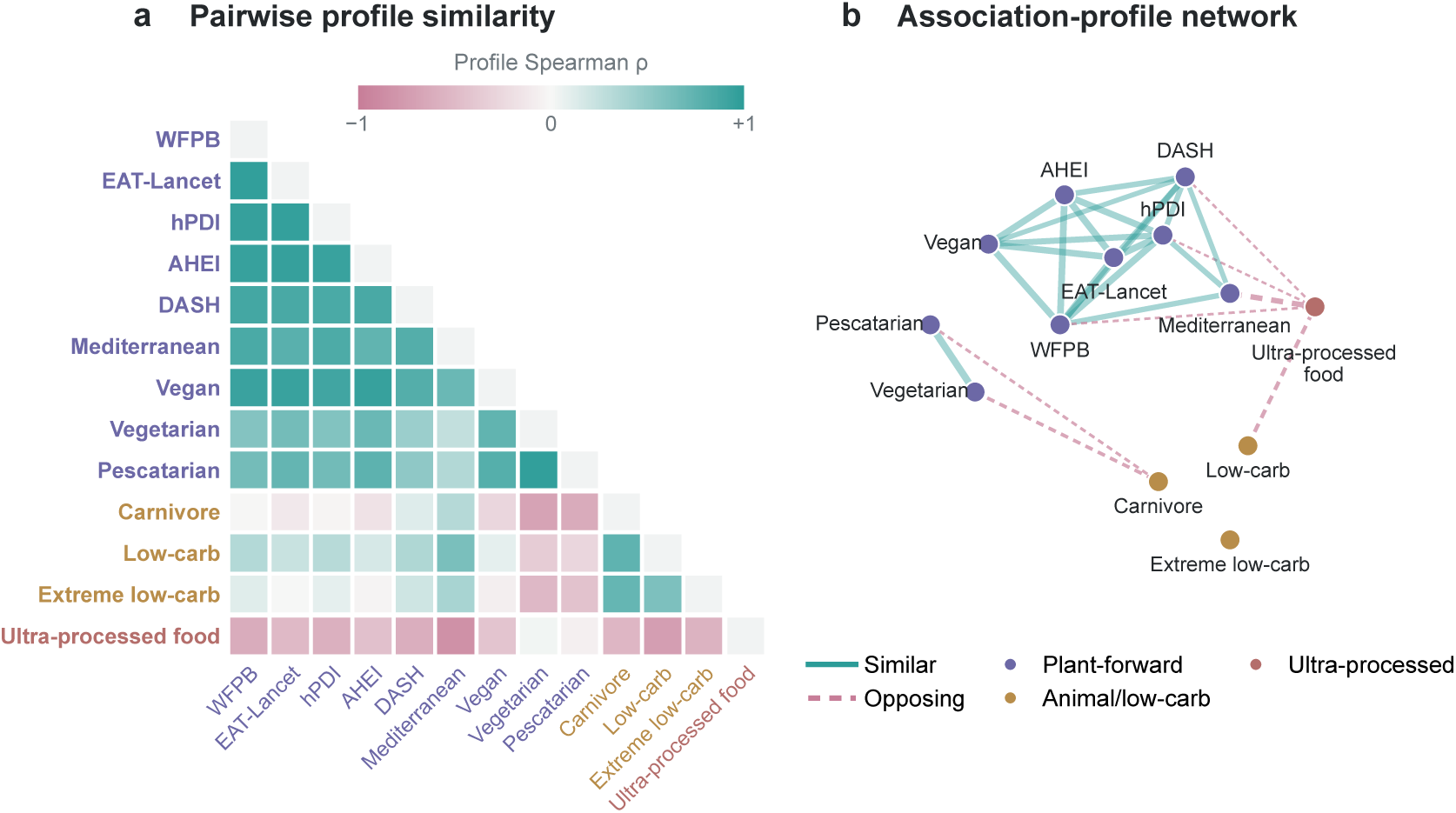
Shared and opposing dietary-pattern association profiles. **a**, Each dietary pattern was represented by its association profile across 136 retained species, and pairwise Spearman correlations were calculated across the 13 patterns. The lower-triangle heatmap encodes the direction and strength of these correlations; teal indicates positive and rose indicates negative correlation. **b**, Network representation of dietary-pattern pairs. Nodes denote dietary patterns and node colours indicate pattern groups; solid teal edges indicate similar profiles and dashed rose edges indicate opposing profiles.

Species within the same genus sometimes showed opposing associations with the same dietary pattern. This occurred in 17 of 120 dietary pattern–genus combinations containing at least two associated species (14.2%). The whole-food plant-based score was inversely associated with most *Streptococcus* species, including *S. gordonii* and *S. oralis*, but positively associated with *S. sanguinis*. Higher overall diet quality has also been linked to lower *S. gordonii* abundance in postmenopausal women [14]. Similar within-genus differences occurred for the carnivore, low-carbohydrate and extremely low-carbohydrate scores in *Prevotella*, *Veillonella* and *Actinomyces*, respectively.

### 2.5 Prediction of oral functional pathways from diet

To examine dietary prediction of functional potential, we assessed whether dietary features improved prediction of oral microbial pathway abundance. After prevalence filtering and quality control, 297 pathway targets were retained. Models including six covariates were compared with models that additionally included 1,400 diet-derived features, using OOF Pearson’s *r*, permutation testing and FDR correction (Fig. 5).

**Fig. 5.**
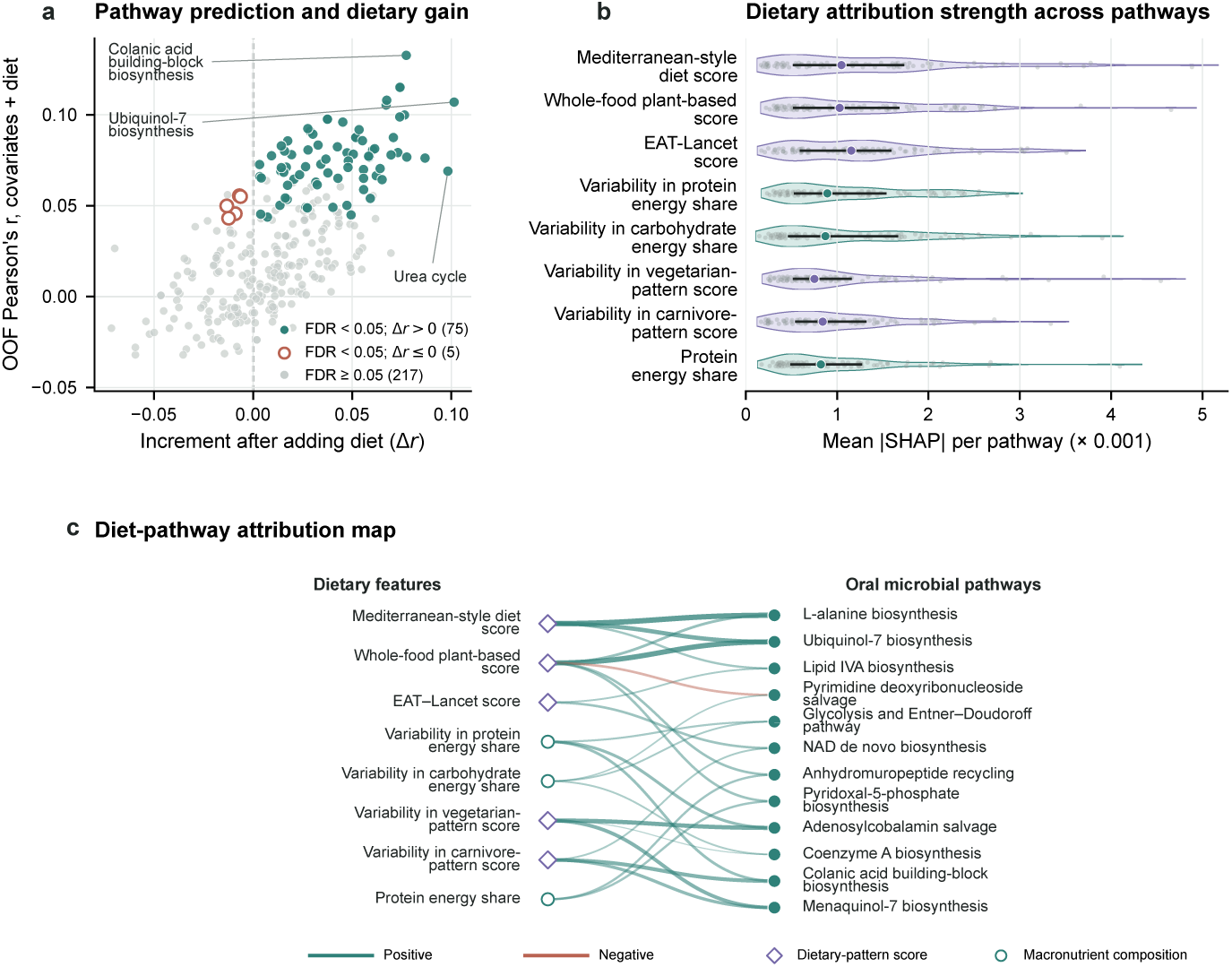
Dietary contributions to oral microbial pathway prediction. **a**, Out-of-fold (OOF) Pearson’s *r* for covariates-plus-diet models (y axis) versus the increment over covariates-only models (Δ*r*, x axis) across 297 pathways. Filled teal and open rust points indicate permutation-FDR-significant pathways (*q <* 0.05) with positive and non-positive Δ*r*, respectively; grey points indicate *q ≥* 0.05. **b**, Mean absolute SHAP distributions for 8 selected dietary features across 80 permutation-FDR-significant pathways. Small points represent pathways; large points and horizontal bars indicate medians and interquartile ranges. **c**, Attribution map for 12 selected pathways with permutation FDR *<* 0.05 and Δ*r >* 0. Each pathway connects to the two features in **b** with its largest mean absolute SHAP values. Edge width indicates mean absolute SHAP; colour indicates the sign of the feature–pathway Spearman correlation. Diamonds and open circles denote dietary-pattern scores and macronutrient-composition features, respectively; filled circles denote pathways.

Adding dietary features modestly improved functional pathway prediction (Fig. 5a). Across 297 pathways, the median Pearson’s *r* increased from 0.0123 in the covariates-only models to 0.0234 in the covariates-plus-diet models, and the mean Pearson’s *r* increased from 0.0155 to 0.0293. Both values were lower than those for species-level abundance. Two-sided Pearson correlation tests between observed abundances and OOF predictions identified 88 significant pathways in the covariates-only models and 123 in the covariates-plus-diet models after FDR correction. Of the 297 pathways, 199 had higher OOF Pearson’s *r* in the covariates-plus-diet models than in the covariates-only models (Fig. 5a).

Covariates-plus-diet models significantly predicted 80 of 297 pathways after permutation-based FDR correction (*q <* 0.05; Fig. 5a; Supplementary Fig. 2). Among the permutation-FDR-significant pathways, colanic acid building-block biosynthesis had the highest observed OOF correlation (COLANSYN-PWY; *r* = 0.133). Other highly ranked pathways included L-alanine, menaquinol, pyridoxal-phosphate and lipid IVA biosynthesis and nucleotide salvage.

We then used SHAP to identify dietary contributors to pathway prediction. Fasting- and eating-window duration and measures of dietary-recording frequency and diversity were prominent predictors. Among the displayed dietary-composition features, Mediterranean-style, whole-food plant-based and EAT–Lancet scores had the highest mean absolute SHAP values averaged across pathways. Protein energy share and day-to-day variability in carbohydrate and protein energy shares and in carnivore and vegetarian scores also contributed (Fig. 5b,c).

Within the cofactor-related pathways, pyridoxal-5*^′^*-phosphate (PLP; vitamin B6) biosynthesis I and both heme b biosynthesis pathways showed weak positive Spearman correlations with Mediterranean-style, whole-food plant-based and EAT–Lancet scores (*ρ* = 0.037–0.058 for PLP and 0.028–0.069 for heme). Flavin and biotin biosynthesis showed inverse correlations with these scores. Among the displayed dietary features, day-to-day variability in vegetarian-pattern score and in protein energy share made the largest contributions to predicting adenosylcobalamin salvage.

### 2.6 Cross-cohort replication in NHANES

We evaluated genus-level replication in 4,667 participants from NHANES 2009–2012 (Fig. 1c). NHANES used one or two 24-hour dietary recalls and oral rinse samples profiled by 16S rRNA gene sequencing. Analyses used 20 shared summary features covering intake amounts, macronutrient composition, eating frequency and timing (Methods 4.7).

To assess whether dietary prediction of oral microbial abundance replicated across cohorts, we fitted genus-level models in NHANES using the 20 shared dietary features (Methods 4.7). The species-to-genus strategy selected candidate genera by mapping significantly predicted HPP species to their corresponding genera. 21 of the 25 evaluable genera met all three replication criteria in NHANES (84.0%). The direct genus-level strategy selected candidates from genus-level prediction analyses in HPP. 12 of the 14 evaluable genera met the same replication criteria in NHANES (85.7%). To assess replication of individual diet–genus associations, we evaluated all 280 combinations of the 20 shared dietary features and 14 evaluable genera in HPP. Of these, 148 pairs met the HPP candidate-selection criteria and formed the fixed set for external replication. In NHANES, 46 of these 148 pairs met the external replication criteria (31.1%). These association-level criteria required an association *q <* 0.05 after correction across all 280 NHANES diet–genus tests, a concordant association direction and improved out-of-fold prediction of genus abundance after adding the corresponding dietary feature.

Most replicated associations involved eating frequency (33 of 46 pairs). In both cohorts, more frequent intake (more daily dietary entries, foods, food categories and one-hour intervals with intake, and longer eating windows) was associated with lower abundances of anaerobic genera, including *Porphyromonas*, *Tannerella*, *Fusobacterium*, *Prevotella*, *Selenomonas* and *Solobacterium*, and with higher *Rothia* abundance. Among nutrient features, dietary fibre intake, per day and per 1,000 kcal, was positively associated with *Lautropia* in both cohorts, consistent with the plant-forward attribution profile of *L. dentalis* in HPP (Fig. 3b). Carbohydrate intake was inversely associated, and protein energy share positively associated, with genera including *Porphyromonas* and *Tannerella*. Together, these analyses support cross-cohort reproducibility of genus-level diet–oral microbiome associations.

## 3 Discussion

By combining detailed dietary records with oral metagenomic data, we identified both shared and species-specific patterns of diet–microbiome association. Several dietary patterns showed similar species-level association profiles, whereas some species within the same genus had opposing associations with a given dietary pattern. Dietary features also provided information for predicting oral microbial composition and functional potential, and some genus-level signals were replicated in the independent NHANES population. Together, these findings connect overall dietary patterns with individual microbial taxa and associations that can be reproduced across populations. Plant-forward and guideline-based scores showed similar association directions and relative strengths across oral species, indicating shared microbial association profiles. Carnivore and low-carbohydrate scores also showed similar species association profiles to one another. The species association profile of ultra-processed food share was negatively correlated with those of most other dietary scores. The HPP gut microbiome study also grouped dietary scores by shared microbial signatures, placing carnivore and low-carbohydrate scores in the same group [23]. These similarities may reflect overlap in the foods and nutrients represented by the scores. A study of three US cohorts found more consistent oral microbiome associations for added sugar than for overall diet quality scores [24]. Integrating dietary-pattern results with food- and nutrient-level findings can help clarify shared microbial features and associations with specific dietary components. Species within the same genus could have opposing associations with a dietary pattern. For example, the whole-food plant-based score was positively associated with *Streptococcus sanguinis* and negatively associated with *S. gordonii* and *S. oralis*. Genus-level summaries may obscure these directional differences among constituent species.

Species composition also provides context for interpreting the diversity findings. Several plant-forward and guideline-based scores were associated with lower oral species richness and Shannon diversity. In postmenopausal women, HEI-2020 was associated with specific oral taxa but showed no significant association with alpha or beta diversity [14]. A separate oral microbiome study found higher diversity in periodontitis than in periodontal health [25]. These observations support interpreting diversity differences alongside the taxa involved and oral health status.

Adding dietary features improved cross-validated prediction of diversity and the abundances of most species and pathways. Contributions from individual foods, dietary-pattern scores and day-to-day variation in intake differed among models predicting diversity, species abundance and pathway abundance. The contribution of eating-time features is consistent with a randomized crossover study in healthy women showing that meal timing altered daily rhythms of salivary microbial diversity [26]. Dietary prediction gains remained positive after adjustment for sampling season in the five species examined.

The functional analyses highlighted dietary associations with the genomic potential for oral microbial cofactor biosynthesis. PLP and heme biosynthesis pathway abundances showed weak positive correlations with Mediterranean-style, whole-food plant-based and EAT–Lancet scores in our cohort. An independent study of Emirati adults reported lower abundances of PLP and heme biosynthesis pathways in participants with obesity than in matched controls [27]. Associations differed across cofactor pathways, indicating pathway-specific relationships with dietary patterns. Transcriptomic and metabolomic measurements could test whether differences in genomic potential correspond to differences in pathway activity or cofactor levels.

In NHANES, 21 of 25 evaluable genera in the species-to-genus strategy and 12 of 14 in the direct genus-level strategy met the replication criteria. These findings identify genus-level diet–microbiome signals that recur across populations, dietary assessments and microbiome measurements. Species-resolved studies could clarify which taxa underlie these replicated genus-level signals.

Absolute prediction correlations remained modest overall across diversity, species and pathway models. Our cross-sectional study characterizes dietary and microbial differences at the same baseline visit. Longitudinal observations are needed to establish the temporal ordering of these associations. The HPP findings reflect the buccal mucosal sampling site, and relative abundances describe compositional relationships among community members. Future studies could prioritize shared dietary-pattern signatures and representative associations with opposing directions among species within the same genus. Repeated sampling and targeted dietary interventions could then examine how these microbial features change with diet.

## 4 Methods

### 4.1 Study population

The Human Phenotype Project (HPP) is an ongoing longitudinal deep-phenotyping cohort of community-dwelling adults in Israel, with recruitment beginning in January 2019. The cohort and its oral metagenomic resource have been described previously [28]. HPP recruitment excluded participants with major acute or unstable systemic diseases, while allowing common stable conditions. Participants reporting oral antibiotic use for gingival or periodontal problems within the previous 3 months were excluded from all oral microbiome analyses (*n* = 10). The present analysis included 5,824 participants with app-based dietary records and oral metagenomic profiles matched to the same baseline visit. Covariate data for the baseline analysis included age, sex, body mass index, smoking status, soft-tissue inflammation level and periodontal damage level. Each participant constituted an independent observational unit.

This research complied with all relevant ethical regulations. The HPP was conducted in accordance with the Declaration of Helsinki and was approved by the Weizmann Institutional Review Board at the Weizmann Institute of Science (reference no. 2392-4). All participants provided written informed consent. At registration, participants self-reported their date of birth and sex. No separate gender variable was collected or used in this study.

### 4.2 Dietary assessment and feature construction

Dietary intake was recorded using the HPP diet-logging smartphone app. Participants were instructed to begin logging one day before an assessment-centre visit and to record everything they ate and drank for at least 14 days after each visit, although the app remained available throughout the study. Each entry recorded the time of consumption, food identity, serving amount, food category and nutritional composition, including energy, macronutrients, sodium, alcohol and dietary fibre. Meals were defined as clock hours containing at least one logged entry.

Dietary records associated with each participant’s baseline visit were processed using the published HPP framework [23]. Foods recorded by fewer than 1% of participants were omitted. Entries were removed when food names or energy values were missing, energy was outside 0–3,000 kcal, or serving weight was outside 0–2,400 g. Foods with incomplete or incorrect nutrient information were excluded, except for sugar substitutes. Duplicate entries were removed. Recording days were excluded if total energy intake was below 500 kcal, above 4,000 kcal or more than 2.5 standard deviations below the participant’s mean daily intake. Participants with fewer than three retained recording days were excluded.

A recording day was defined as a calendar date containing at least one retained dietary entry. Mean daily energy intake from each food or food category was calculated by dividing its total recorded energy by the participant’s number of recording days.

Dietary features characterized food and nutrient intake, dietary patterns, dietary variety and eating timing. Additional measures captured the duration and frequency of dietary recording. Food-level and food-category features captured daily intake and the percentage of energy contributed by individual items or categories. Nutrient features included daily intake, intake per 1,000 kcal and macronutrient energy percentages.

Standard dietary scores included the Israeli Mediterranean Diet Adherence Screener (IMEDAS), EAT–Lancet index and Dietary Approaches to Stop Hypertension (DASH) score. We also calculated the Alternative Healthy Eating Index (AHEI) and healthful Plant-Based Diet Index (hPDI). The hPDI assigned higher scores to healthy plant foods and lower scores to less healthy plant foods and animal foods.

Food-alignment scores quantified the fraction of daily energy supplied by foods eligible for each dietary pattern. The whole-food plant-based (WFPB) score included unprocessed plant foods, whereas the vegan score also included processed vegan foods. The vegetarian score additionally allowed dairy products and eggs, and the pescatarian score also allowed fish and seafood. The carnivore score included animal-derived foods. For patterns with intake thresholds, energy shortfalls in required food groups and excesses in restricted groups were subtracted from eligible-food energy before division by total daily energy. For WFPB, daily minimums were 100 kcal each from vegetables and fruit, and 5% of total energy from each of three groups: whole grains, pulses, and nuts and seeds. For the carnivore score, energy from hard cheese, milk, cream cheese and yoghurt above 50% of total daily energy incurred a penalty. Food-alignment scores were bounded between 0 and 1.

Ultra-processed food intake was expressed as the percentage of daily energy supplied by foods classified as NOVA group 4. Low-carbohydrate and extremely low-carbohydrate scores represented the proportions of retained recording days with carbohydrate energy shares of 10–26% and below 10%, respectively.

Daily dietary features, including daily adherence scores, were summarized by participant-level means and sample standard deviations. For selected features, coefficients of variation were calculated as the sample standard deviation divided by the participant-level mean.

In total, the feature matrix contained approximately 1,400 diet-derived variables. A complete feature dictionary, including units and derivation formulae, will be provided in the Supplementary Information.

### 4.3 Oral microbiome profiling and outcome construction

Oral samples were collected at baseline HPP visits using the standardized bilateral buccal-mucosa swab protocol described previously [28]. Participants were asked to avoid tooth brushing or antiseptic mouthwash for at least 4 h and eating for at least 1 h before sampling; water was allowed. Immediately before sampling, participants rinsed once with water and waited 10–15 min. Trained staff wearing sterile gloves used single-use flocked swabs to rub and rotate along the left and right buccal mucosa with moderate pressure. Contact with the teeth, gingival crevices, tongue dorsum and saliva pools was avoided. The resulting profiles primarily represent the buccal mucosal community. Swabs were placed into collection tubes immediately, frozen at *−*80 *^◦^*C within 2 h and shipped on dry ice for downstream processing. Collection and handling conditions are summarized in Table 2.

**Table 2.** Buccal-swab collection and handling conditions.

| Stage | Collection and handling requirements |
| --- | --- |
| Participant preparation | Avoid tooth brushing and antiseptic mouthwash for $\geq 4$ h, and food intake for $\geq 1$ h; water is permitted. |
| Before collection | Rinse once with water, followed by a 10–15-min interval before sampling. |
| Specimen collection | Trained staff wearing sterile gloves collect samples from both buccal surfaces using single-use flocked swabs, with rubbing and rotation under moderate pressure. |
| Sampling boundaries | Avoid the teeth, gingival crevices, tongue dorsum and pooled saliva. |
| Sample handling | Transfer swabs immediately into collection tubes, freeze at $-80^{\circ}\text{C}$ within 2 h, and transport on dry ice. |

DNA extraction and library preparation were performed at Gencove Laboratories. Whole-genome sequencing was performed at Neogen GeneSeek Laboratories on an Illumina NovaSeq platform using 150-bp paired-end reads. Raw reads were aligned to the human reference genome, and non-aligned reads were retained for microbial analysis. Sequencing adapters and low-quality sequences were removed using Trimmomatic [29]. Taxonomic composition was profiled from quality-controlled non-human reads using MetaPhlAn 4 [30], which generated species-level relative abundances. Functional potential was quantified using HUMAnN 3.6, generating gene-family and pathway abundance and coverage profiles [28, 31].

We analysed four classes of oral microbial outcomes: species richness, Shannon diversity, species-level relative abundance and pathway abundance. Species richness was defined as the number of species detected in each sample before zero replacement. Shannon diversity was calculated from species-level relative abundances as *H* = − Σ*_f_ p_f_* ln(*p_f_*), where *p_f_* is the relative abundance of species *f* within a sample.

Species and pathways detected in at least 200 participants were retained, following the previously described HPP oral metagenomic pipeline [28]. For each retained feature, zero values were replaced with half the first percentile of its non-zero abundances across participants. Within each participant, abundances were then divided by their sum across retained features, scaled to parts per million and log_10_-transformed. Species and pathway profiles were processed separately.

### 4.4 Dietary pattern association analyses

We used covariate-adjusted linear models to assess associations of all 13 dietary pattern scores with oral microbial species abundance, species richness and Shannon diversity. The primary model adjusted for age, sex, body mass index and smoking status; the extended model additionally adjusted for soft-tissue inflammation level and periodontal damage level.

Within each adjustment model, Benjamini–Hochberg correction was applied across all dietary pattern–species tests and, separately, across the 26 dietary pattern– alpha-diversity tests. Associations with an FDR-adjusted *q <* 0.05 were considered significant. We summarized the number and direction of associations significant in both models.

For each dietary pattern, its species association profile comprised primary-model regression coefficients (*β*) across all retained species. Profile similarity was quantified using Spearman’s correlation. To examine opposing associations within genera, we grouped associations significant in both models by dietary pattern and genus. Groups containing at least two associated species were classified as having opposing directions when their primary-model *β* coefficients included both positive and negative values.

### 4.5 Prediction of oral microbiome features from diet

We assessed whether adding dietary features improved prediction of oral microbial diversity, species abundance and pathway abundance. For each microbial target, we fitted a covariates-only model and a covariates-plus-diet model. Both models included the six prespecified covariates described above and used gradient-boosted decision trees implemented with the LightGBM regressor [32]. The same deterministic five-fold cross-validation splits, generated without shuffling, were used for both models. Predictions for the held-out participants from all five folds were concatenated to obtain one out-of-fold (OOF) prediction vector per target.

Age and body mass index were treated as continuous variables. Sex, smoking status and the two oral-health measures were represented using their HPP assessment values. For model fitting, any remaining missing predictor values were imputed using training-fold medians, and continuous predictors were standardized using training-fold means and standard deviations. Binary predictors were not standardized. The target was standardized within the training fold before model fitting, and held-out predictions were transformed back to the original target scale before evaluation. These transformations were applied to the corresponding held-out fold without recalculation. LightGBM hyperparameters were fixed across targets: regression objective, maximum tree depth of 4, 2,000 boosting iterations, learning rate of 0.001, row subsampling fraction of 0.5 with a subsampling frequency of 1, column subsampling fraction of 0.3, *L*_1_ and *L*_2_ regularization coefficients of 1, and a minimum of 20 observations per leaf. The model random seed was 1. Predictive performance was quantified using Pearson’s correlation coefficient between observed and OOF-predicted values and the coefficient of determination (*R*^2^). Incremental predictive information from diet was summarized as Δ*r* = *r*covariates+diet *− r*covariates only.

For pathway models, two-sided Pearson correlation *P* values were adjusted across all retained pathways using the Benjamini–Hochberg procedure, separately for each model type.

The significance of the cross-validated predictions of the covariates-plus-diet models was assessed by permutation testing. Alpha-diversity outcomes were tested individually. Species and pathways were grouped into 20 prevalence bins of 5% width (0–5%, 5–10%, …, 95–100%), and the null distribution obtained for one representative target in each bin was applied to all targets in that bin. In each of 500 permutations, target values were randomly reassigned across participants, with predictors and cross-validation folds held fixed, and the complete fold-specific preprocessing and modelling procedure was repeated. One-sided permutation *P* values were calculated as

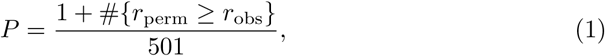

where *r*_obs_ is the observed OOF Pearson’s *r* and *r*_perm_ are the values from the permuted datasets. *P* values were adjusted using the Benjamini–Hochberg procedure separately within the alpha-diversity, species-abundance and pathway-abundance analysis layers, and targets with *q <* 0.05 were considered significantly predicted.

For the five species shown in Fig. 3b, we assessed sensitivity to oral sampling season using the same participants as in the primary analysis. Seasons were defined from sampling month as spring (March–May), summer (June–August), autumn (September–November) and winter (December–February, reference). Season indicators were added to both models, with preprocessing, hyperparameters and five-fold splits unchanged.

### 4.6 Interpretation of model predictions using SHAP

We used SHapley Additive exPlanations (SHAP) [33] to interpret the fitted covariates-plus-diet models. Feature importance for a target was summarized as the mean absolute SHAP value across participants. SHAP analyses were performed for both alpha-diversity outcomes and for species and pathways that passed permutation-based false-discovery-rate correction. For pathway models, SHAP values were computed for a subset of 1,000 participants on the standardized target scale.

For visualization, directional SHAP summaries were calculated as mean absolute SHAP values multiplied by the sign of the feature–target Spearman correlation. Signs indicate feature–target association directions rather than raw signed SHAP effects. The fitted models and complete attribution outputs included host covariates and record-derived logging and eating-pattern variables. Selected main-figure displays showed dietary-composition features.

### 4.7 External replication in NHANES

We used data from the 2009–2010 and 2011–2012 cycles of the US National Health and Nutrition Examination Survey (NHANES) to independently replicate the diet-related oral microbiome signals identified in HPP. After matching dietary, oral microbiome and covariate data, we retained participants aged 30–69 years with complete data for all six covariates. The analysis included 4,667 participants, comprising 2,134 males (45.7%) and 2,533 females (54.3%), with a mean age of 49.2 years. The NHANES oral microbiome data were derived from oral rinse samples [34]. The V4 region of the 16S rRNA gene was amplified using the 515F and 806R primers and sequenced on the Illumina HiSeq 2500 platform. Sequences were denoised and processed for amplicon sequence variant inference using DADA2 [35], with taxonomic assignment based on SILVA v123.

HPP used continuous app-based dietary records, whereas NHANES provided up to two 24-hour dietary recalls per participant. Only 24-hour dietary recalls classified as reliable by NHANES were retained, with each recall day assessed separately. Dietary features were calculated separately for each valid recall day and then averaged within participants. Nutrient densities and energy percentages were calculated at the daily level before averaging.

We harmonized dietary variables across cohorts by nutritional meaning, calculation formula and measurement unit, retaining 20 shared dietary features. These comprised total energy intake; the daily numbers of dietary entries, distinct foods and top-level Food Patterns Equivalents Database (FPED) food categories; the number of one-hour clock intervals containing at least one dietary entry; the daily number of distinct foods divided by the number of these intervals; daily eating-window duration; mean daily intakes of carbohydrate, fat, protein, dietary fibre, sodium and alcohol; dietary fibre and sodium intake per 1,000 kcal; the percentages of energy derived from carbohydrate, protein, fat and alcohol; and the proportion of valid recall days on which carbohydrate provided 10–26% of total energy.

All NHANES prediction and association models included the same six covariates: age, sex, BMI, current smoking status, self-reported gum disease and a history of non-traumatic tooth mobility. The two oral-health measures provided approximate counterparts to the HPP inflammation and periodontal-damage measures, although their questions and response scales differed.

We selected candidate genera through two complementary strategies: species-to-genus mapping and direct HPP genus-level prediction analyses.

The species-to-genus strategy selected candidates by mapping significantly predicted HPP species to their corresponding genera. The 101 retained HPP species that passed permutation-based FDR correction yielded 42 candidate genera, of which 25 could be matched to NHANES. For these genera, we compared covariates-only models with models additionally including the 20 shared dietary features.

The direct genus-level strategy selected candidates from HPP genus-level prediction analyses using the complete dietary feature matrix. We selected genera with FDR-significant covariates-plus-diet predictions. Their dietary prediction increments also had to pass permutation-based FDR correction. Of the 22 HPP candidate genera, 17 had exact genus-name matches in the two released sequence-filtering versions. Eligibility additionally required detection in at least 200 participants and non-zero variance in log_10_-transformed abundance in the primary dataset. These criteria yielded 14 genera, which were retained for the six-covariate replication analysis.

NHANES prediction analyses used the same training-fold preprocessing, cross-validation scheme and model settings as the HPP discovery cohort (Methods 4.5). For both genus-level replication strategies, prediction significance was assessed using two-sided Pearson correlation tests between observed abundances and OOF predictions. Benjamini–Hochberg correction was applied across the evaluated genera, separately for each model and replication strategy.

Dietary prediction increments in both genus-level strategies were evaluated by jointly permuting the 20-feature dietary block within each NHANES cycle while keeping outcomes, covariates and cross-validation folds fixed.

Genus-level replication was determined in the primary NHANES analysis. It required a covariates-plus-diet model *q <* 0.05, an observed Δ*r >* 0 and an increment-test *q <* 0.05. Model prediction and dietary increment tests were corrected separately across the genera evaluated within each replication strategy.

We also evaluated replication of individual diet–genus associations across the 14 evaluable genera and 20 shared dietary features. For each pair, we tested whether adding that dietary feature to the covariates-only model improved OOF prediction of genus abundance. In NHANES, covariate-adjusted ordinary least squares regression estimated association effect sizes and directions using log_10_-transformed genus abundance. Dietary features entered the regression in their original units, and coefficient significance was assessed using conventional standard errors and two-sided *t* tests. Both association and prediction models used the same six covariates and participants. Association *P* values were adjusted across all tested diet–genus pairs in HPP using the Benjamini–Hochberg procedure. Pairs with association *q <* 0.05, a non-zero regression coefficient and Δ*r >* 0 formed a fixed set of 148 HPP candidates. All 280 diet–genus combinations were also evaluated in NHANES, with association *P* values adjusted across all 280 tests. Replication was assessed within the fixed 148-pair HPP candidate set and required an association *q <* 0.05, concordant regression-coefficient signs between cohorts and a positive OOF Δ*r* after adding the dietary feature.

## Data availability

The Human Phenotype Project (HPP) data used in this study are available through the HPP data-access framework. Raw oral metagenomic sequencing data and basic participant phenotypes are available through the European Genome-phenome Archive under accession number EGAS50000001763, where applicable. Other participant-level HPP data used in this study, including app-based dietary records, oral-health variables and linked phenotype data, are subject to controlled access because of privacy and ethical restrictions and can be requested through the HPP data-access portal (https://humanphenotypeproject.org/data-access). Bona fide researchers should contact for access instructions.

NHANES dietary, demographic, smoking and oral-health variables are available from the public NHANES data resources. Processed oral microbiome abundance data used for the replication analyses are available from the corresponding NHANES data release. Participant-level raw oral microbiome sequencing data are restricted and must be accessed through the NCHS Research Data Center application process. Associated sequencing quality-control data are available under NCBI SRA BioProject accession PRJNA896165.

## Code availability

Code used to construct the dietary features, perform the HPP prediction and association analyses, harmonize dietary variables across cohorts, and perform the NHANES replication analyses is available at the GitHub repository https://github.com/haochen-MBZUAI/Diet-Oral-Microbiome.

## Author contributions

Kaiyu Liang performed the data analysis, developed the methods and implemented the visualizations. Huifa Li and Yulong Li assisted with data processing. Haochen Xue, Anastasia Godneva and Imran Razzak wrote the manuscript. Jionglong Su, Xiaojun Zhu, Sibei Chen and Yichen Li reviewed and revised the manuscript. Imran Razzak and Eran Segal conceived and supervised the project.

## Appendix

**Supplementary Fig. 1.**
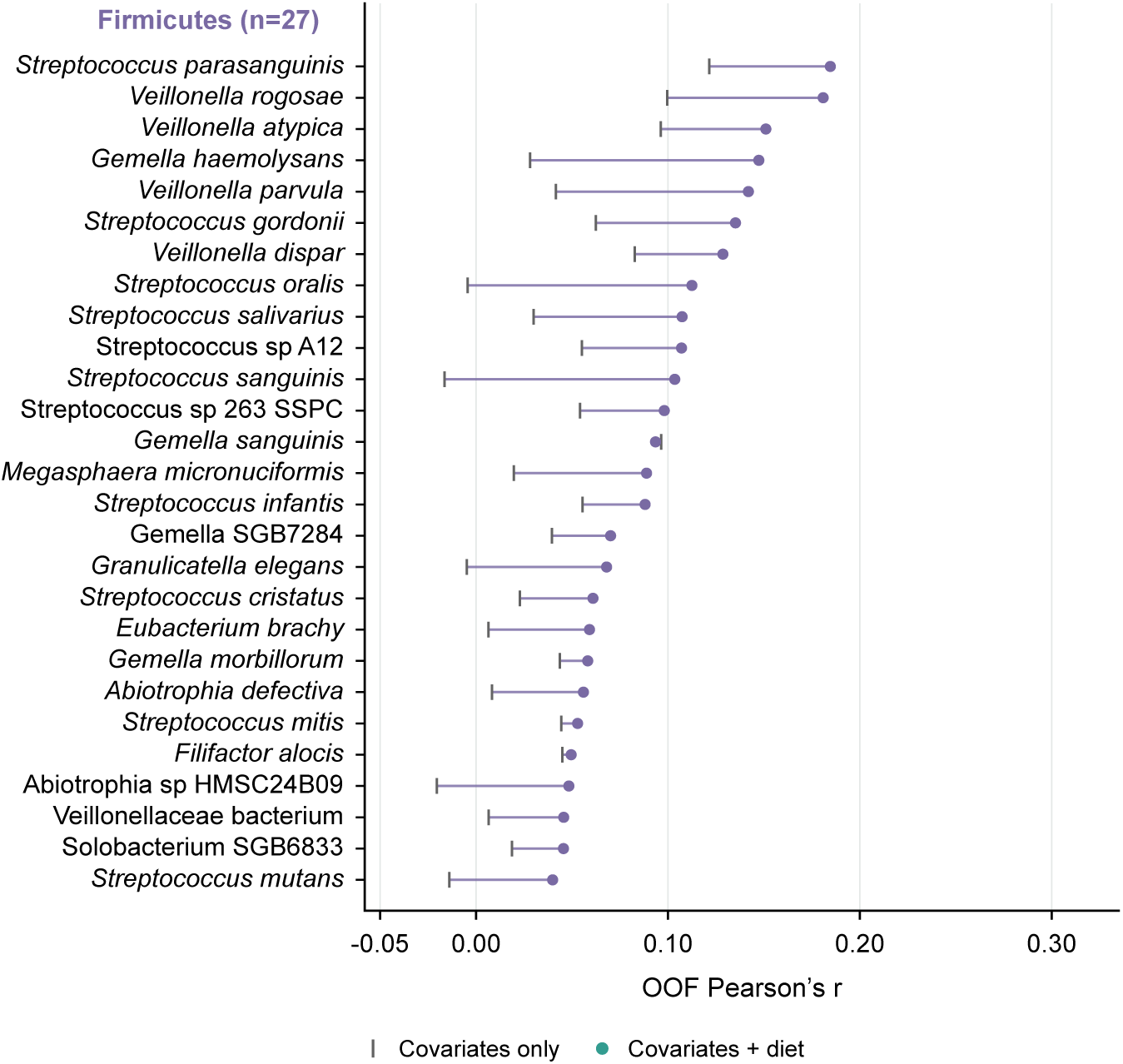
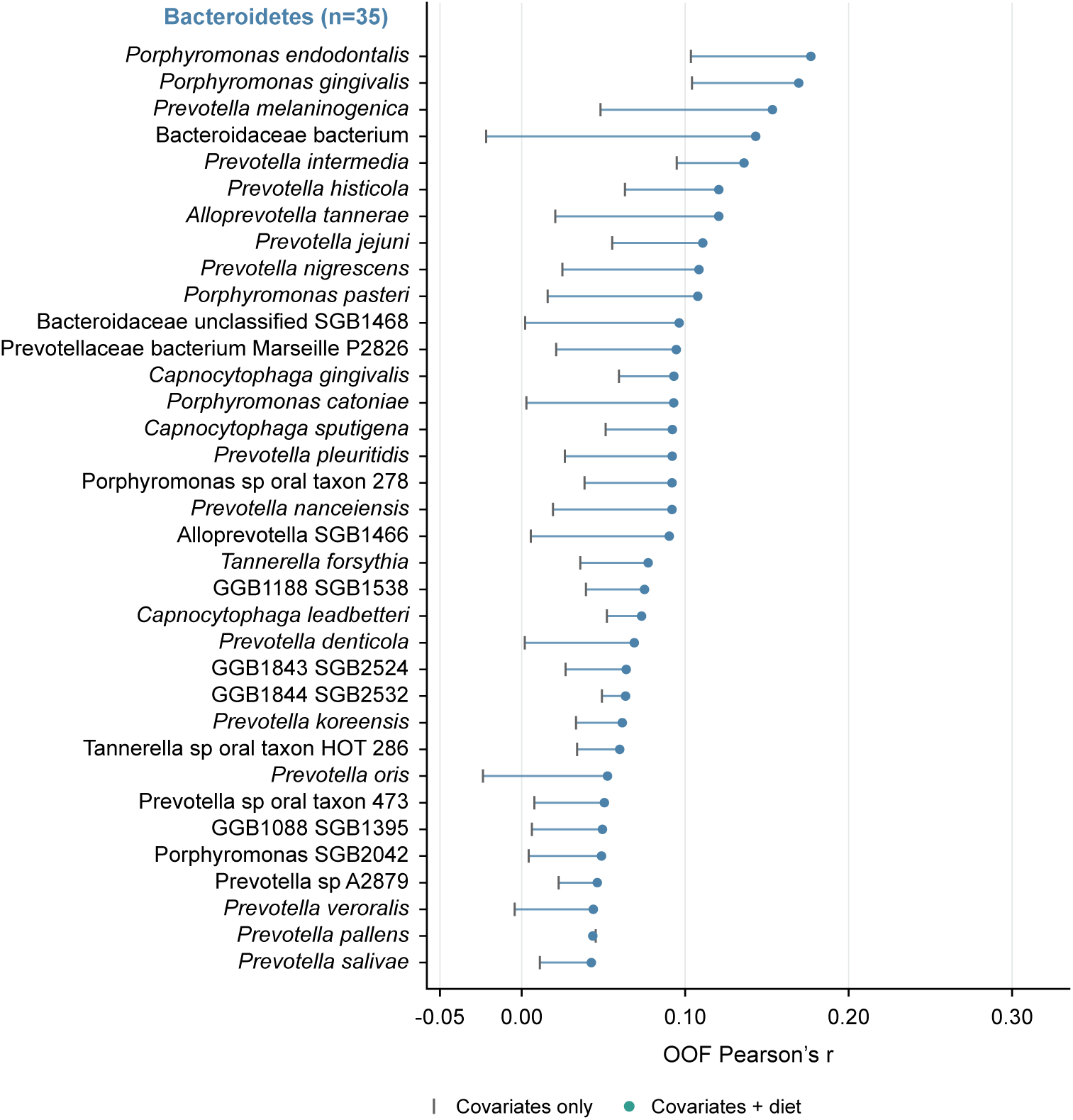
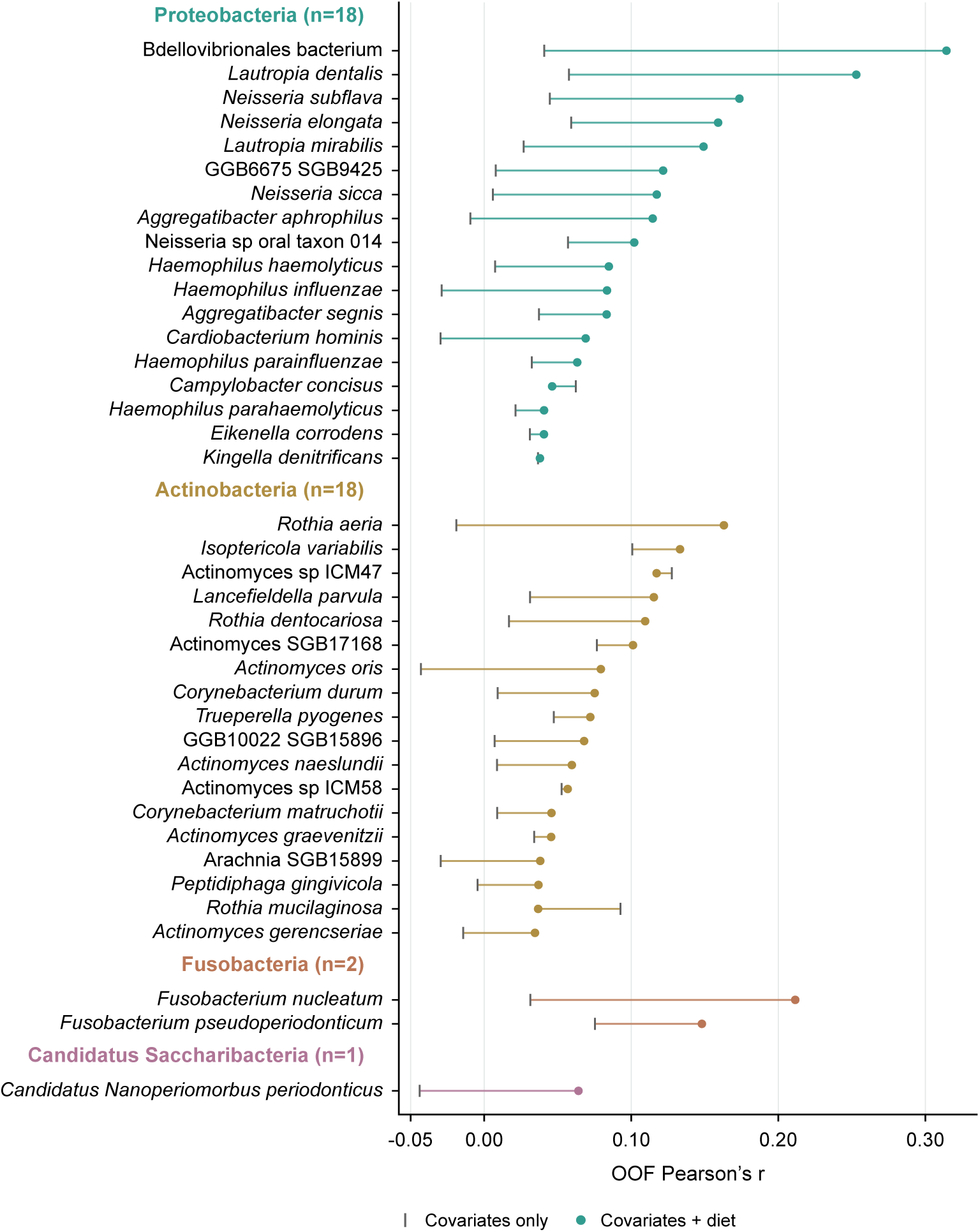
Prediction performance of individual oral species. OOF Pearson’s *r* for covariates-only models (grey vertical ticks) and covariates-plus-diet models (coloured circles) for the 101 species with permutation-FDR-significant covariates-plus-diet predictions (*q <* 0.05). Lines connect the two model performances for each species. Species are grouped by phylum and ordered by covariates-plus-diet *r* within each group.

**Supplementary Fig. 2.**
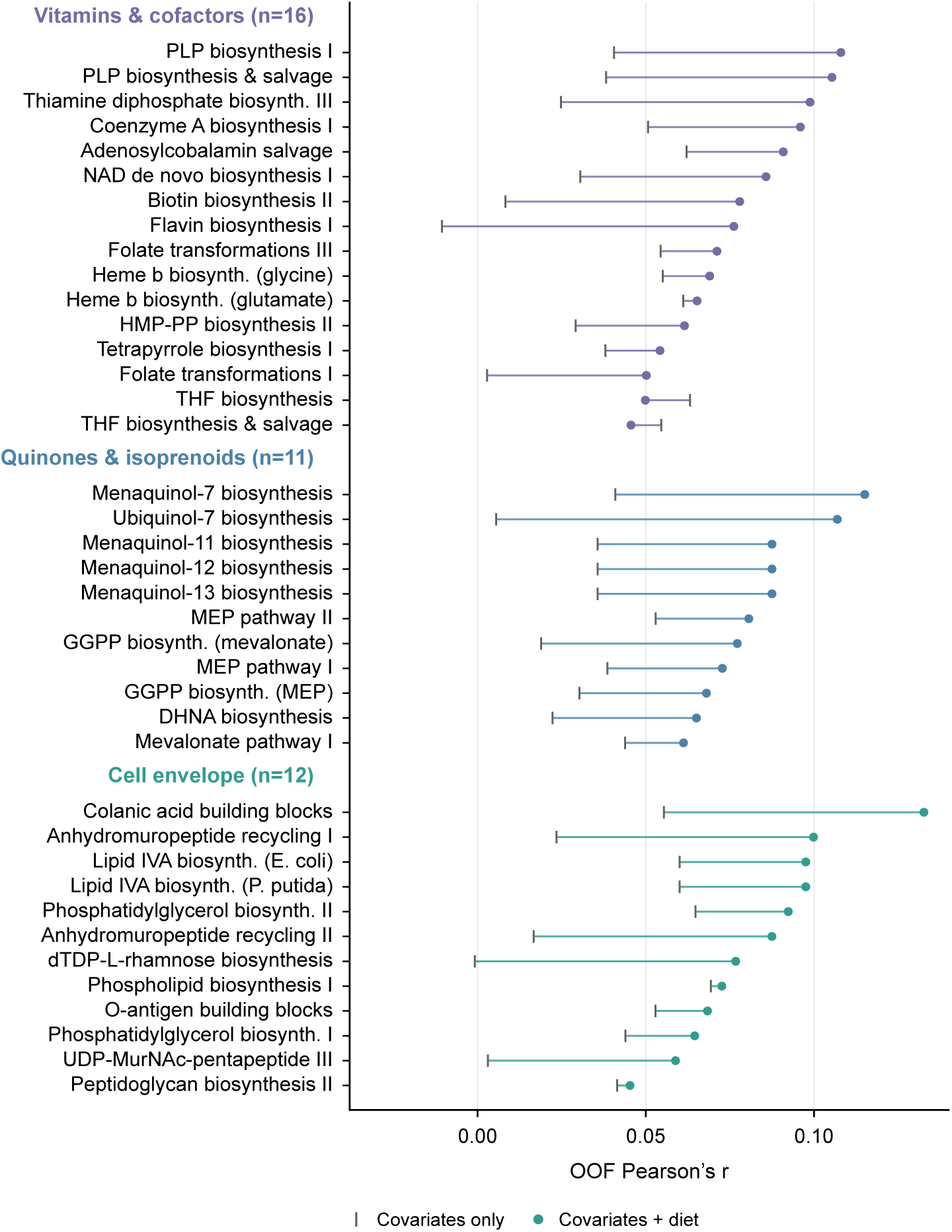
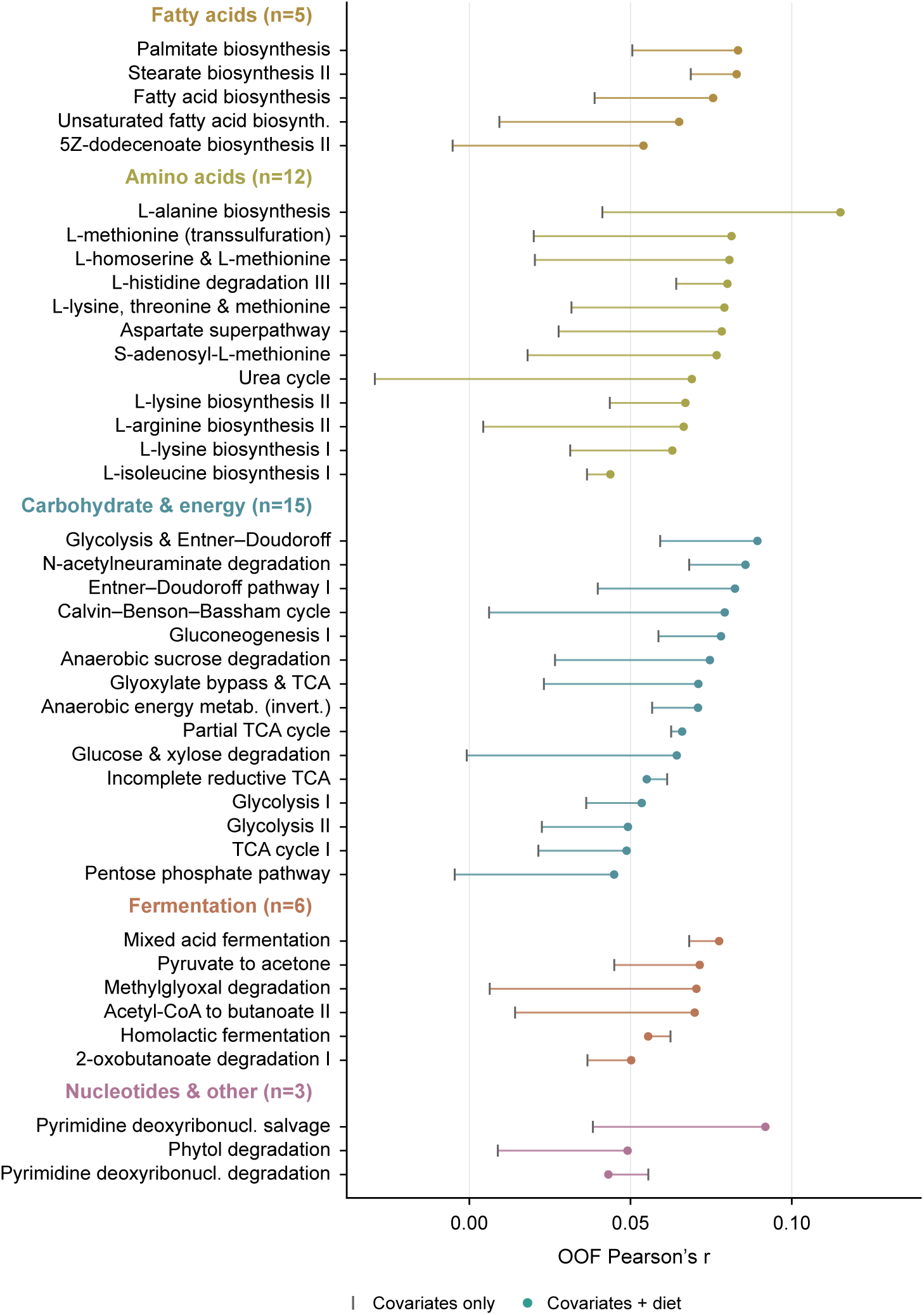
Prediction performance of individual oral microbial pathways. OOF Pearson’s *r* for covariates-only models (grey vertical ticks) and covariates-plus-diet models (coloured circles) for the 80 pathways with permutation-FDR-significant covariates-plus-diet predictions (*q <* 0.05). Lines connect the two model performances for each pathway.

